# Oscillatory brain states govern spontaneous fMRI network dynamics

**DOI:** 10.1101/393389

**Authors:** Daniel Gutierrez-Barragan, M. Albert Basson, Stefano Panzeri, Alessandro Gozzi

## Abstract

Spontaneous brain activity as assessed with resting-state fMRI exhibits rich spatiotemporal structure. However, the principles by which brain-wide patterns of spontaneous fMRI activity reconfigure and interact with each other, remain unclear. We devised a frame-wise clustering approach to map spatiotemporal dynamics of spontaneous fMRI activity with voxel resolution in the resting mouse brain. We show that brain-wide patterns of fMRI co-activation can be reliably mapped at the group and subject level, defining a restricted set of recurring brain states characterized by rich network structure. We document that these functional states are characterized by contrasting patterns of spontaneous fMRI activity and exhibit coupled oscillatory dynamics, with each state occurring at specific phases of global fMRI signal fluctuations. Finally, we show that autism-associated genetic alterations result in the engagement of non-canonical brain states and altered coupled oscillatory dynamics. Our approach reveals a new set of fundamental principles guiding the spatiotemporal organization of resting state fMRI activity, and its disruption in brain disorders.

## Introduction

Spontaneous neural activity is ubiquitously present in the mammalian brain and persists across physiological states (Buzsáki and Draguhn, 2004; Mohajerani et al., 2010). In humans, whole-brain patterns of intrinsic brain activity are typically mapped by measuring spontaneous functional magnetic resonance imaging (fMRI) signal in the resting brain (Power et al., 2014), an approach termed resting-state fMRI (rsfMRI). A large body of experimental work has shown that low-frequency fluctuations in the fMRI signal are temporally synchronous across multiple functional systems, delineating a set of reproducible topographies known as resting-state networks, which can be reliably identified also in primates (Vincent et al., 2007) and rodents (Gozzi and Schwarz, 2016). These observations have prompted a widespread use of inter-regional correlation between rsfMRI signals as an index of functional coupling, or “functional connectivity” between regions. Importantly, additional studies have further shown that rsfMRI activity is characterized by rich temporal structure, involving dynamic reconfiguration into transient states entailing variations in resting state networks occurring on the time scale of seconds (Hutchison et al., 2013a). These studies have promoted a view of the resting brain as an inherently dynamic system, in which highly evolving patterns of instantaneous activity interact over time in a sporadic or stochastic fashion (Braun et al., 2015).

Recent animal studies have linked hemodynamic-based measures of intrinsic brain activity to low-frequency oscillatory neural activity as measured with calcium imaging (Matsui et al., 2016; Schwalm et al., 2017; Xiao et al., 2017). Specifically, optical imaging studies in mice have implicated global waves of neural activity and transient neural co-activations among homotopic areas as key neural drivers of hemodynamic-based measurements of functional connectivity (Matsui et al., 2016, 2018). In keeping with these experimental results, human rsfMRI network activity can be reliably described by brief instances of regional peak fMRI activity (Liu and Duyn, 2013), a feature that has been related to transient variation in calcium co-activation patterns (Matsui et al., 2017). Taken together, these findings suggest that reconfiguration of spontaneous network activity may be guided by transitions between recurring patterns of slow-wave activity. Such an interpretative framework is consistent with the initial recognition of putative temporal sequences of propagated fMRI activity defined as quasi-periodic patterns, or lag threads of propagated fMRI signal in human rsfMRI datasets (Mitra et al., 2015; Yousefi et al., 2018). However, a precise characterization of how network states interact and dynamic brain reconfigurations occur is lacking. Are spontaneous network transitions organized stochastically, or do they reflect a set of recurring default states? And what are the fundamental principles by which brain-wide patterns of spontaneous fMRI activity reconfigure and interact with each other?

Here we devised an optimized frame-wise clustering strategy, combining the identification of recurrent patterns of spontaneous fMRI activity with an analysis of how these spatial patterns are dynamically coupled. We show that this approach enables brain-wide mapping of spontaneous rsfMRI activity with voxel-resolution in the resting mouse brain (Sforazzini et al., 2014; Liska et al., 2015; Gozzi and Schwarz, 2016), and demonstrate the reproducibility of our findings across different rsfMRI datasets. Specifically, we describe a set of recurring brain-wide functional states that can be reliably mapped at the group and subject level, and show that the dynamics of their transitions can be compactly described as a set of coupled oscillators, each corresponding to a spatial pattern, and each preferentially occurring at specific phases of global fMRI signal fluctuations. Importantly, we demonstrate that aberrant patterns of fMRI connectivity in a genetic model of autism reflect the engagement non-canonical brain states, characterized by altered regional topography and oscillatory dynamics. Collectively, our approach points at oscillatory network activity as a fundamental level of organization of spontaneous brain activity, and describes a set of new principles guiding the spatio-temporal organization of resting state activity.

## Results

### Selective fMRI frame averaging recapitulates networks of correlated activity

The observation that spontaneous brain activity may be driven by brief instances of simultaneous activation of various brain regions (Matsui et al., 2016), implies that key information about the dynamic structure of resting state activity can be retrieved from individual fMRI volumes (here on referred to as ‘frames’). It has been previously shown that selective averaging of fMRI frames exhibiting *regional* peaks of BOLD activity can closely recapitulate rsfMRI connectivity networks obtained via seed-based correlation analysis (Tagliazucchi et al., 2012; Liu and Duyn, 2013). As a first step towards a voxel-wise mapping of spontaneous fMRI signal dynamics in the mouse, we probed whether this relationship holds true also in this species. To this aim, we measured rsfMRI network activity in 40 adult male mice over a time window of 10 minutes (500 timepoints, main dataset). We used pre-defined anatomical regions as rsfMRI correlation seeds, and spatially averaged individual fMRI frames as a function of their peak intensity signal, to produce seed-based mean co-activation patterns (CAPs). We next compared the obtained “seed-based mean CAPs” with canonical rsfMRI correlation maps (Fig.1). Consistent with previous observations, we found that previously-described rsfMRI networks, including the mouse hippocampal, latero-cortical, auditory-temporal and default-mode networks (DMN) (Liska et al., 2015), can be spatially reproduced by averaging a limited number (e.g. 15%, Fig. 1A) of fMRI frames exhibiting peak BOLD activity in corresponding anatomical seed locations.

**Figure 1.**
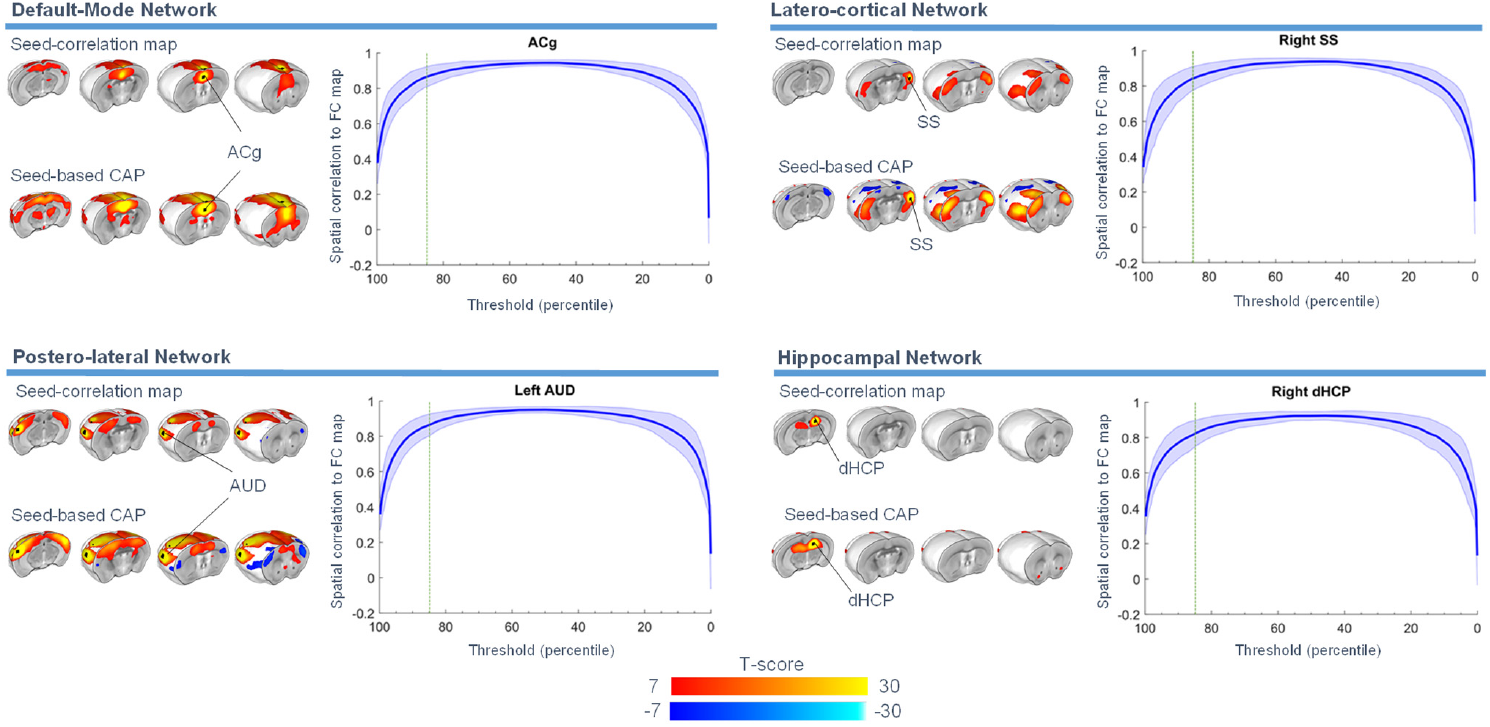
Selective fMRI frame averaging recapitulates rsfMRI network activity. rsfMRI networks obtained via seed-based correlation are spatially recapitulated by averaging fMRI frames exhibiting peak regional BOLD fMRI activity (seed-based CAPs). This relationship is illustrated for four representative mouse rsfMRI networks mapped via seed-based correlation analysis (ACg, anterior cingulate, SS, somatosensory cortex, AUD, auditory cortex, dHCP, dorsal hippocampus). Plots on the right illustrate the spatial overlap between seed-based correlation maps and seed-based CAPs, as a function of the percentage of frames used for the computation of the latter (group mean +/− SEM). The dashed green line indicates the 15- percentile threshold employed for seed-based CAP visualization.

### Whole-brain fMRI frame clustering reveals a set of recurring functional brain states

The observation that regional peaks of fMRI activity drive spatially-structured network topographies is consistent with spontaneous neural activity being a non-stationary phenomenon, in which evolving brain-states undergo recurring reconfiguration. To obtain a regionally-unbiased characterization of these states and their transition dynamics in the mouse brain, we devised a k-means clustering analysis of all the individual rsfMRI frames, without any *a priori* temporal, anatomical or intensity-based restriction (Fig. 2). As opposed to describing co-activated networks as a set of spatially-correlated patterns of activity, this approach identifies, as prototypes of each of *k* spatial activity clusters, the different types of simultaneous single time-frame co-activation patterns (shortened hereafter as CAPs or “functional states”) that recur in the data (Fig 2C). In contrast to seed-guided selective fMRI frame averaging, this strategy does not require the *a priori* selection of regions of interest or the use of fMRI intensity thresholds, and can identify composite functional states characterized by both patterns of simultaneous co-activation (above fMRI signal baseline) and co-deactivation (below fMRI signal baseline, Fig. 2). Importantly, because our approach classifies activity in all time frames, it permits to reliably detect multiple brain-wide patterns of spontaneous brain activity, and not only those that occur at either local or global peaks of regional BOLD activity.

**Figure 2.**
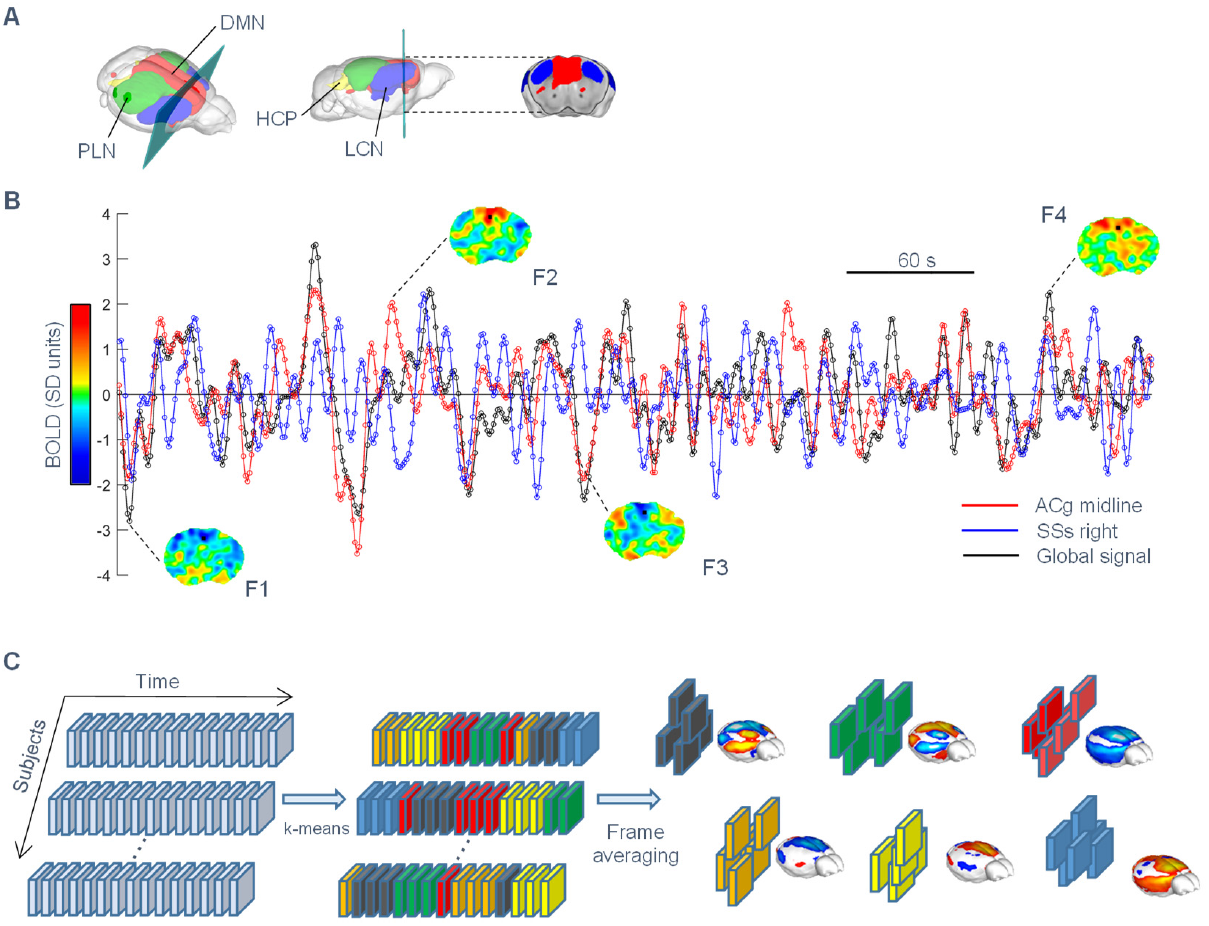
Identification of recurring brain states via whole-brain fMRI frame clustering. (A) Glass brain representation of the seed-based resting-state networks described in Figure 1 (DMN: default mode network, HCN: hippocampal network, LCN: latero-cortical network, PLN: postero-lateral network) (B) Illustrative fMRI BOLD time course (SD units) in the anterior cingulate (ACg, red), somatosensory cortex (SSs, blue) as well as fMRI global-signal (black) in a representative subject (brain slice illustrated in panel A). Note the presence of peaks of concordant or diverging BOLD activity in cingulate and somatosensory areas across time, suggestive of time-varying network reconfiguration; cingulate and somatosensory regions are concurrently co-activated and co-deactivated in F1 and F4, but they exhibit opposing BOLD activity in F2 and F3. (C) These dynamic transitions can be captured and classified into recurring brain states by clustering fMRI frames into spatially congruent patterns (CAPs), using the k-means algorithm.

The use of an unsupervised algorithm like k means poses the problem of selecting an appropriate number of clusters. Our goal was to partition the dataset into whole-brain fMRI states that are robust and reproducible, and to ensure that each selected cluster could be trusted to reflect a genuine set of coactive brain regions. To this aim, we first computed, for increasing k, how much variance is explained by the clustering algorithm (Fig. S1B). The explained variance curve, computed for the main dataset, revealed an elbow region encompassing the range k = 4 – 10 in which variance was still increasing (thus using more cluster yielded a better description of the datasets), but its increase was progressively smaller (denoting that the importance of adding more clusters was getting smaller and smaller, Fig. S1, B and C). We selected the value k as the highest value within the elbow region that still ensured maximal between-dataset reproducibility of the corresponding CAPs with respect to two additional independently collected rsfMRI datasets (n = 41 and n = 23, respectively). With this procedure, we identified k = 6 states which as robustly conserved across the three rsfMRI datasets, and necessary for describing the datasets with high accuracy (Fig. 3 and Fig. S1). Further credibility for the robustness of these six states as sets of genuinely coactive fMRI voxels, is given by the identification of the same CAPs at the single subject level (described below), and the fact that these 6 patterns were present also when partitioning the main dataset into a higher number of clusters (Fig. S1). While fMRI clustering with k =7 and 8 revealed additional plausible states, we restricted our subsequent analyses to the first 6 clusters, as this appears to be the finest datasets partition exhibiting the highest cross-dataset reproducibility. It should however be noted that there was a seventh states, plotted in Fig. S2, that appeared to be very stable across the two larger wild-type datasets (n = 40 and n = 41), but that exhibited low reproducibility on the third, smaller, reference image set (n = 23). Although we did not include this seventh state in any further analyses here, our observations raise the possibility that this CAP, being found very robustly in the two largest independent datasets, may reflect a less stable, yet functionally meaningful state, and so we briefly document its properties in the supplemental information section (Supplemental text).

**Figure 3.**
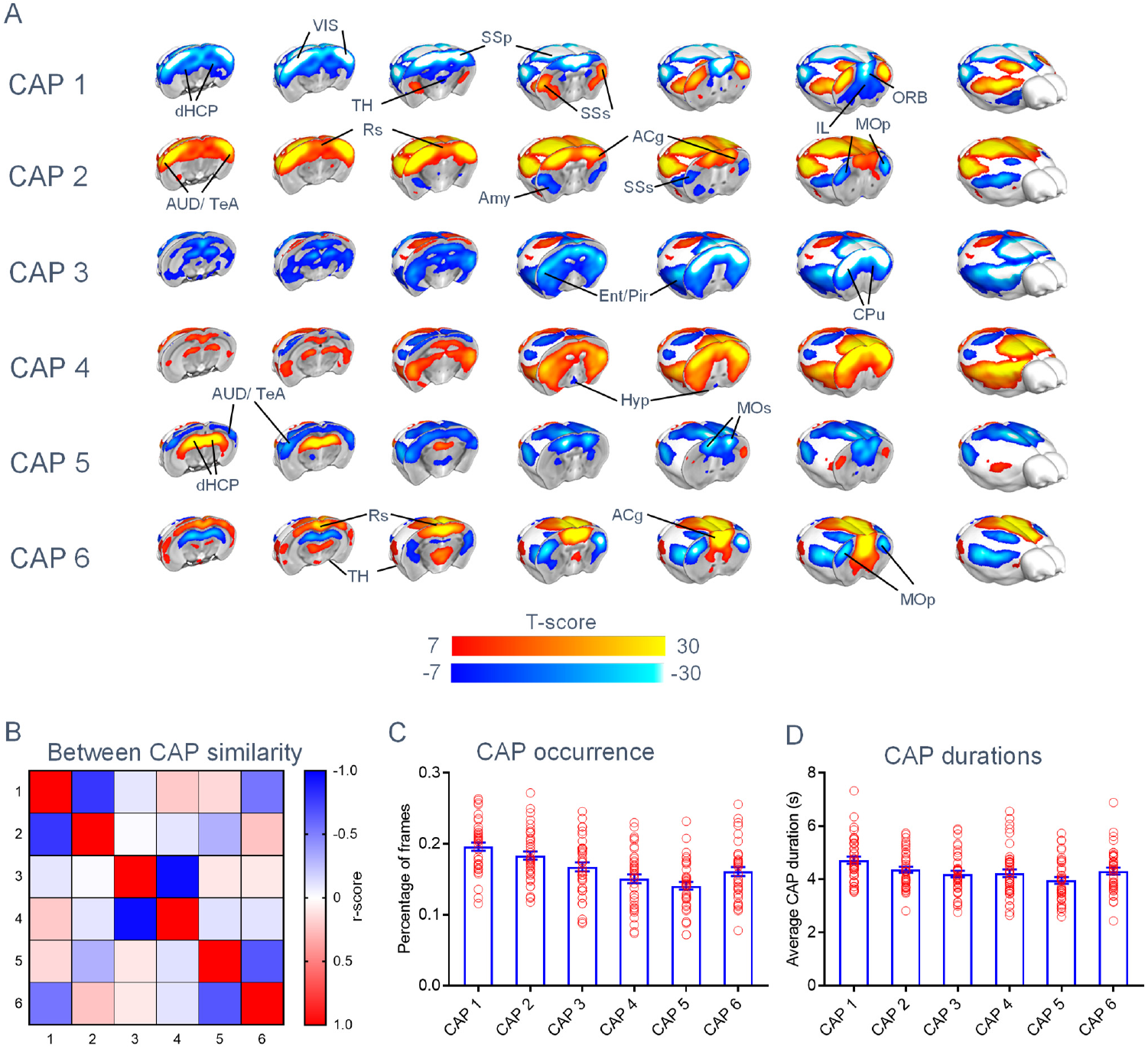
Recurring functional states of the mouse brain. (A) Whole brain representation of the functional brain states (CAPs) we identified at the group level. Red/yellow indicates co-activation (i.e. high fMRI BOLD signal) while blue indicates co-deactivation (i.e. low fMRI BOLD signal) (p < 0.01, Bonferroni corrected). CAPs have been ordered based on their spatial properties by numbering consecutively states characterized by opposing BOLD co-activation patterns (i.e. 1-2, 3-4 and 5-6), as denoted by the negative correlations in panel (B). Panels (C) and (D) illustrate CAP occurrence rate and mean duration, respectively (mean +/− SEM). Abbreviations: ACg – Anterior Cingulate cortex; AUD – Auditory cortex; dHC – dorsal Hippocampus; vHC – ventral Hippocampus; HT – Hypothalamus; ILA – Infralimbic Area; LAN – Lateral Amygdalar Nucleus; MOp – primary Motor cortex; Mos – secondary Motor cortex; ORB – Orbitofrontal cortex; PIR – Piriform Area; PL – Pallidum; Rs – Retrosplenial cortex; SSp – primary Somatosensory cortex; SSs – secondary Somatosensory cortex; ST – Striatum; TeA – Temporal Association cortex; TH – Thalamus; VIS – Visual cortex.

### fMRI states encompass known connectivity networks of the mouse brain and can be identified at the single subject level

An illustration of the six identified states is reported in Fig. 3A. One defining characteristic of all the six CAPs is their configuration as a composite assembly of regional substrates encompassing previously-described distributed resting-state networks of the mouse brain. For example, CAP1 shows a clear co-activation of primary and secondary motor-sensory areas belonging to the mouse latero-cortical network (LCN) together with deactivation of cortico-limbic regions and peri-hippocampal constituents of the mouse DMN. Similarly, CAP 5 encompasses co-deactivation of the DMN, and co-activation of the hippocampal network. These correspondences are illustrated in Fig. S3, in which we report an empirical decomposition of some of these CAPs into a set of putative constituting rsfMRI networks. The presence of spatially-prominent contributions of known rsfMRI connectivity networks in the identified states, plus their duration in the in the order of a few seconds (Fig. 3D), implicate the observed patterns as time-varying functional states underlying rsfMRI network dynamics as assessed with correlational techniques.

To investigate whether the selected six states are representative of spontaneous brain dynamics identifiable at the single subject level, we repeated our state-detection using k = 6 on a subset of nineteen mice for which we acquired rsfMRI images over a 30-minute window. We next spatially matched each subject-level state with the corresponding CAP obtained at the group-level analyses (Fig. 3A), to obtain a voxel-wise CAP incidence maps. These analyses revealed that all 6 states can be reliably identified at the single subject level, with foci of very high cross-subject incidence in key network locations (Fig. 4A). Importantly, the spatial distribution of the observed CAPs at the subject level (Fig. 4B) closely recapitulates the features that we observed with group-level clustering (Fig. 4B). These correspondences suggest that the six identified CAPs correspond to genuine brain states, representative of spontaneous functional reconfigurations occurring at the single subject level.

**Figure 4.**
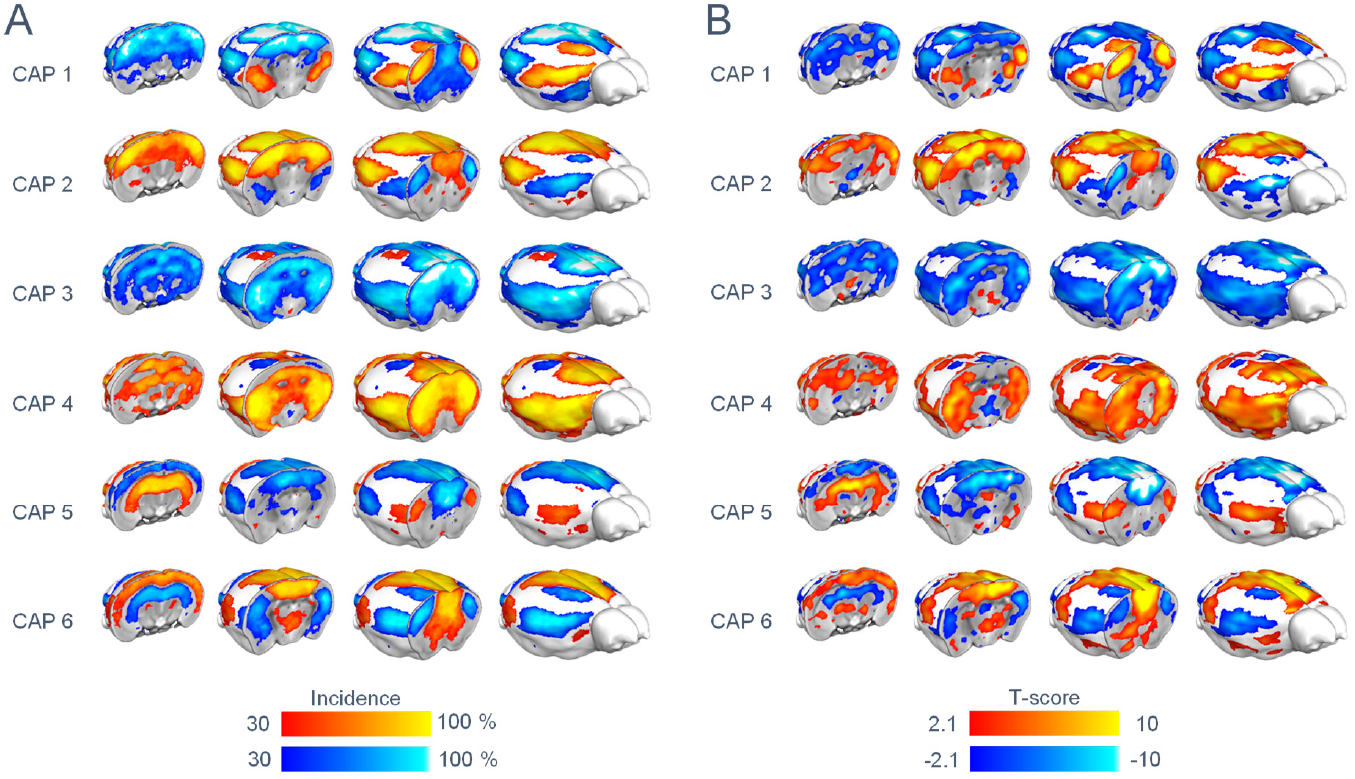
Brain states can be detected at the single-subject level. Left: incidence map of each CAP at the subject level (k = 6, 30 min rsfMRI acquisitions, p < 0.05, FDR corrected). Each voxel represents the proportion of subjects with significant co-activation (p < 0.05, FDR corrected) as its corresponding group-level CAP template. Right: Spatial distribution of CAPs detected in a representative subject (p < 0.05, FDR corrected). Yellow indicates regional co-activation, blue regional co-deactivation.

The use of sedation in mouse fMRI acquisitions allows for tight control of motion-related artefacts (Gozzi and Schwarz, 2016). We nevertheless assessed the role of potential frame displacements in our datasets by re-computing group-level CAPs upon strict censoring of putative “motion-affected” frames. To this purpose, we employed a frame-wise displacement (FD) threshold of 75 and 100 µm, leading to a rejection of 24% and 10% putative “high-motion” frames, respectively. Despite the use of disproportionately strict motion censoring, the resulting CAPs were spatially undistinguishable from what observed by using uncensored frames (Fig. S4A-B). We also computed the assignment of putative “motion-affected” frames to each CAP using the above-mentioned FD thresholds, and did not observe any CAP being dominantly enriched in motion contaminated frames (one-way ANOVA, p = 0.51 and 0.91 respectively, Fig. S4C-D). These results argue against a significant contribution of motion artefacts to our imaging results.

### Functional states can be classified into opposing, oscillating patterns of co-activation

A notable feature of the identified CAPs, is their configuration into state and anti-state pairs characterized by opposing patterns of functional co-activation (Fig. 3A-B). This feature was especially prominent in CAPs 3 and 4 (r = −0.96), involving a contrasting co-activation of neocortical regions, but was also apparent in CAPs 1 and 2 (r = −0.80), and CAPs 5 and 6 (r = −0.66), the first pair being characterized by a clear anti-correlation between the DMN and LCN, the latter between hippocampal areas and the DMN (Fig. 3A-B). Importantly, the observation of opposite spatial configurations was not necessarily expected, or resulting from the employed clustering procedure. This attribute suggests that the networks expressed by these states continue to be similarly correlated or anticorrelated with other elements of the network not only when these elements are co-activated, but also when they are co-deactivated. This could happen if the state anti-state pairs undergo an ongoing oscillation, with opposite states reflecting peaks and troughs of these network fluctuations.

Based on the above results, we hypothesized that the dynamics of transition between CAPs could be described in oscillatory terms. To test this hypothesis, we computed at each instant of time the spatial correlation between each CAP and the BOLD fMRI signal in that time frame (Fig. 5A). The obtained index, hereafter referred to as “CAP time course”, assesses the spatial match between spontaneous brain activity and the specified CAP in the considered time frame. When we computed the power spectra of each CAP time course (Figure 5B), we observed a clear peak of power in the 0.01-0.03 Hz frequency band, indicating that that brain-wide spontaneous brain activity undergoes transitions between network configurations described by CAPs, with infra-slow oscillatory dynamics. To capture and visualize the spatio-temporal dynamics of CAP assembly and disassembly across consecutive frames, we next computed the average of the whole-brain BOLD frames time-locked around each CAP time course’s local maxima. These results, which are best visualized in the form of movies (Supplementary movies 1-6), clearly show how each state builds-up and disassembles with spatial patterns that strikingly resemble wave-like propagating activity observed with calcium imaging in the mouse dorsal cortex (Matsui et al., 2017).

**Figure 5.**
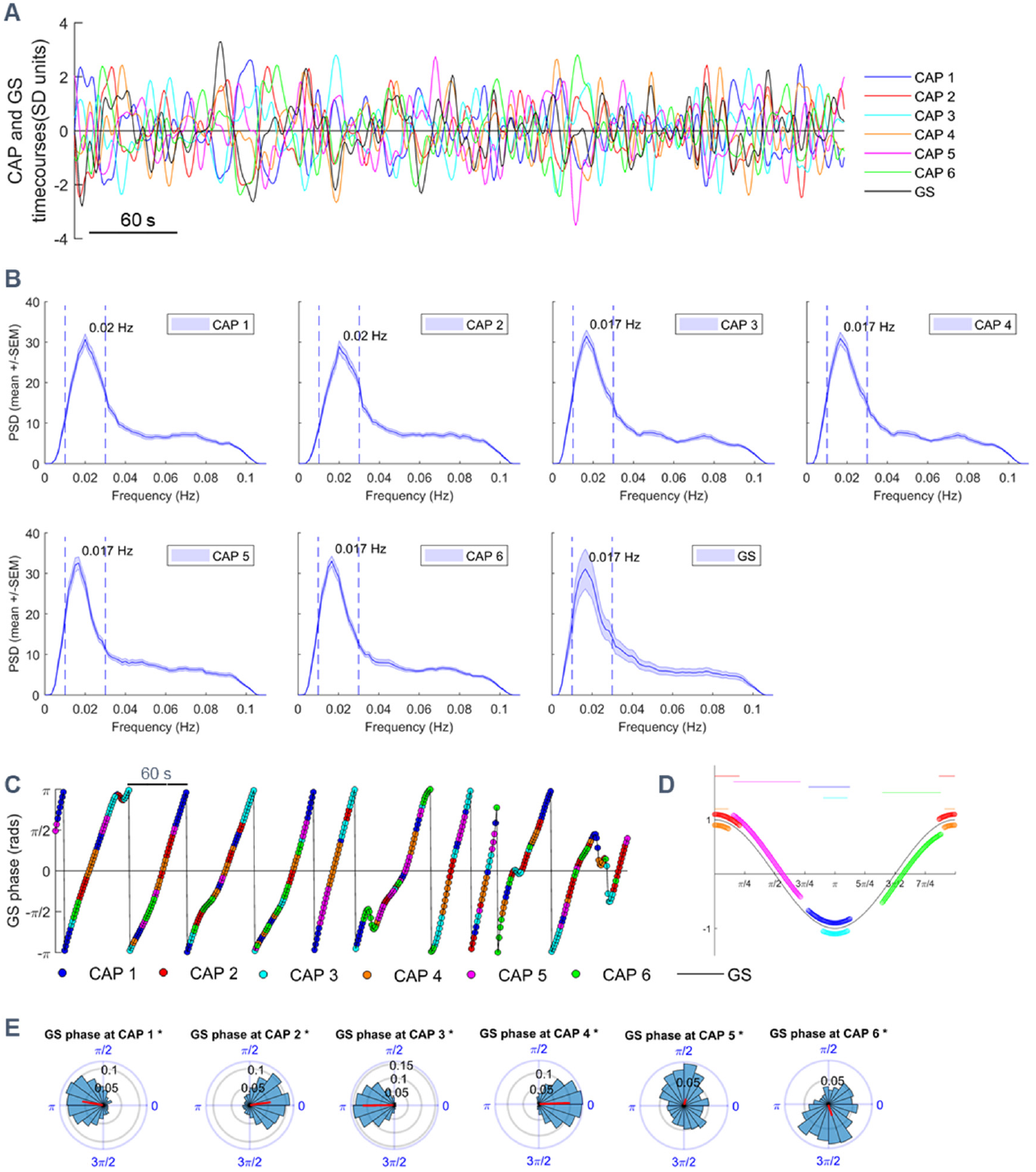
Functional states exhibit oscillatory dynamics. **(A)** Illustrative CAP (or the GS) timecourse from in a representative subject. **(B)** Mean power spectral density of CAPs and the GS (mean +/− SEM). Blue dashed vertical lines delimit the 0.01-0.03 Hz frequency band employed in subsequent phase analyses. (C) Instantaneous phase of the GS in a representative subject. Colored dots mark the occurrence of each CAP. (D) CAPs occur at specific phases of global signal oscillations. Phase dispersion plotted as circular variance around the circular mean (top horizontal bars). CAP encoding is described according to color scheme in (C). E) Circular distribution of GS phases at each CAP’s occurrence within a GS cycle. For each distribution, the resulting vector (magnitude and phase) is shown as a black radial line.

### Functional states transitions occur at specific phases of global fMRI signal

Interestingly, our power analyses also revealed that the fMRI global signal (GS) presents a power spectrum peak in the same infra-slow oscillatory band that characterizes CAP oscillations (Fig 5B). This finding raises the question of how CAP oscillatory dynamics may relate to that of the GS. We hypothesized that intrinsic oscillations in GS may not reflect global, spatially undifferentiated, ups and downs of whole-brain activity, but, on the contrary, each phase of the GS may encompass the specific activation of selected subsets of possible brain states, each characterized by a characteristic profile of brain activity. To test this hypothesis, we measured the frequency of CAP occurrence at different phases of the fMRI GS oscillations, by filtering the GS signal in the infra-slow band and computing the phase at each instant (Montemurro et al., 2008). With the phase convention we used, phase values of 0 and pi corresponded to peaks and troughs of the GS, respectively (Fig. 5D). To understand whether different sections of GS oscillation cycles correspond to different states, we next computed the circular distribution of GS phases at which each CAP occurred. Notably, we found that the occurrences of all 6 CAPs were not distributed uniformly across the GS infra-slow cycle (Fig. 5C-D). Rather, CAP occurrences were concentrated at specific ranges of the GS phase cycle, with all distributions exhibiting a significant deviation from circular uniformity (Raleigh test, p < 0.05, Bonferroni corrected). Specifically, CAPs pairs 3 and 4 as well as 1 and 2 tended to occur around the trough and the peak of GS fluctuations, respectively (Fig. 5D-E), albeit with different spread of phases, while occurrences of CAPs 5 and 6 were concentrated around time points in which the GS phase was 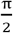 and 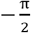 respectively, corresponding to intermediate, off-peak GS levels. The observation that off-peak GS values correspond to a well-defined spatial structure lends strong support to our hypothesis that different phases of the GS oscillations do not reflect just spatially-unstructured ups and downs of global brain activity. The qualitative differences highlighted above were confirmed by quantitative tests, with evidence of significant differences between GS mean phases of all CAPs (Watson-Williams test, p < 0.05, Bonferroni corrected), with the only exception of CAPs 2 and 4.

### Functional states act as coupled oscillators

Although the above analysis suggests that CAPs act as oscillating networks, it does not fully clarify whether they act independently, or as coupled oscillators that covary according to specific phase relationships between each other, and with respect to the global fMRI signal. If the latter is the case, phase relationships between CAPs must be observed not only when pooling all phase occurrences together (as in Fig. 5E), but also when considering phase relationships between occurrences of different CAPs within the same (or immediately adjacent) GS cycle. We thus computed the GS phase angular difference between CAP occurrences in the same GS cycle or across GS cycles that were immediately adjacent in time (i.e. next or previous GS cycle, Fig. S5). We found that GS phase difference between occurrences of the same CAP (diagonal panels, Fig. S5) were concentrated around zero, suggesting that a given state appears in general for a short range of adjacent phases during a given cycle. Moreover, reciprocal CAPs appeared with a phase difference of π in the same cycle, suggesting that each GS cycle reflects at least in part the alternation between peaks and troughs of a specific spatially-structured network. Finally, we found significantly locked distribution of phase differences between non-reciprocal CAP pairs (e.g. between CAPs 4 and 5, and CAPs 3 and 6). Collectively, these results show that the identified brain-wide functional states act like coupled infra-slow oscillators, and suggest that GS fluctuations, rather than being the result of unstructured changes of activity, reflect coupled dynamics of interacting structured networks along the entire oscillation cycle.

### Altered patterns of rsfMRI connectivity entail non-canonical functional state dynamics

The relationship between patterns of spontaneous fMRI activity and correlational rsfMRI network topographies (Fig. 1) offers the opportunity to reframe aberrant rsfMRI connectivity in terms of altered spatial-temporal structure of instantaneous fMRI states. As an illustrative example of this novel interpretative framework, we mapped functional states in mice haploinsufficient for the chromatin remodeling gene *Chd8*. This mutation recapitulates a major genetic risk factor for autism spectrum disorders (ASD), and is characterized by rsfMRI over-connectivity between hippocampal and motor cortical areas, as assessed with conventional steady-state rsfMRI mapping (Suetterlin et al., 2018).

As described above, brain state identification using k = 6 in control animals (*Chd8*^+/+^; n = 23) recapitulated the states observed in our two larger control datasets (Fig. S1). However, corresponding CAPs in Chd8 mutants (*Chd8*^+/−^; n = 21) were characterized by several notable non-canonical spatial features, entailing dysfunctional or aberrant engagement of specific regional substrates (Fig. 6B, T(40) > 2.8, cluster corrected, p < 0.05). Specifically, CAPs 1 and 2 in *Chd8*^+/−^ mice exhibited an aberrant reciprocal involvement of thalamic and dorsal hippocampal regions. Similarly, cingulate and mid-thalamic recruitment appeared to be defective in CAPs 3 and 4 of *Chd8*^+/−^ mice and foci of aberrant motor sensory and prefrontal co-activation were found in CAPs 5 and 6. These results provide evidence of non-canonical state recruitment in mice harboring a key ASD-risk mutation.

**Figure 6.**
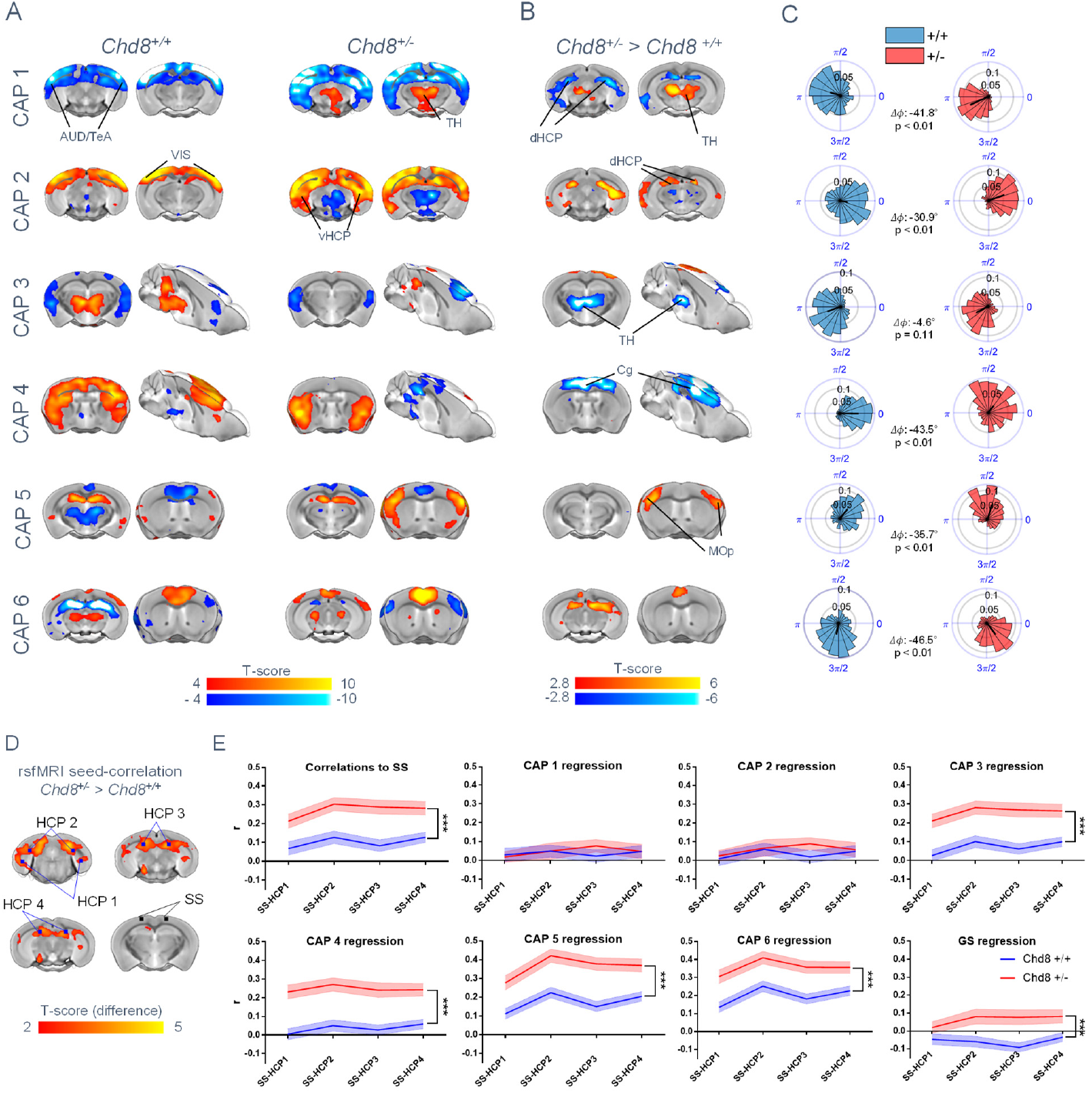
Altered brain states in a genetic mouse model of autism. (A-B) CAPs exhibit non-canonical patterns of co-activation and de-activation in *Chd8*^+/−^ mutants (p < 0.05, FWE cluster corrected) (C) Delayed CAP occurrence within GS cycles with respect to control mice in *Chd8*^+/−^ mutants (p < 0.01, William-Watson test for circular mean homogeneity, Bonferroni corrected, all CAPs, except for CAP3). (D). Seed-based rsfMRI correlation differences in Chd8 mutants. Inter-group differences (p < 0.05, FWE cluster corrected) are depicted with respect to a seed-pair in the somatosensory cortex (SS) to replicate the rsfMRI findings reported in Suetterlin et al., 2018. The location of hippocampal (HPC) seed used for correlational profiling is also illustrated (HPC1, 2 and 3) (E) Seed-based correlation profiling upon regression of individual CAP time courses or the GS in both groups (***p < 0.001, two-way repeated measures ANOVA, mean ± SEM). Abbreviations: Cg – Cingulate cortex; AUD – Auditory cortex; HCP – Hippocampus; dHCP – Dorsal hippocampus; vHCP – ventral hippocampus; MOp – primary Motor cortex; SS – somatosensory cortex< TH – Thalamus.

We next investigated the presence of between-group alterations in CAP oscillatory phase dynamics (Fig. 6C), by mapping the occurrence of each states with respect to GS phase in control and *Chd8* mutants. Interestingly, we found that, although the CAPs of *Chd8*^+/−^ mutants happened along the phase cycle in the same order as the corresponding state of control mice, there were significant genotype-dependent differences in CAP dynamics. Specifically, all the functional states of *Chd8*^+/−^ mice exhibited a significantly delayed occurrence (−30.9°< *Δφ* <−46.5°) with respect to GS oscillation phases (Fig. 6C, p < 0.01, William-Watson test), with the exception of CAP 3, in which no genotype-dependent phase delay was observed. Similarly, the distribution of phases happening within the same or adjacent GS cycles (Fig. S6) showed that all such phase distributions significantly deviate from circular uniformity in both mutant and control mice (Raleigh test, p < 0.05, Bonferroni corrected), showing that CAPs evolve as coupled oscillators in both cohorts. However, there was a significant genotype-dependent difference in the circular mean of the phase distributions of functional states occurring within same or adjacent cycles, with the only exception of phase relationships between CAPs 1-4 and 2-4 (Williams-Watson test, p < 0.05, Bonferroni corrected), with major GS phase differences occurring between CAP 3 and CAPs 4-6. Collectively, these findings suggest that ASD risk mutations can alter rsfMRI connectivity via the recruitment of non-canonical brain states, and by altering coupled oscillatory dynamics of individual brain states.

To better relate dysfunctional state dynamics to aberrant rsfMRI connectivity as assessed with conventional correlation-based measurements, we next investigated whether specific non-canonical states could account for the increased cortico-hippocampal rsfMRI coupling previously described in *Chd8*^+/−^ mice (Suetterlin et al., 2018). To this purpose, we anatomically profiled rsfMRI correlations between the affected somatosensory and hippocampal areas before and after regressing each CAP’s time-course independently (Fig. 6E). Notably, we observed that independent regression of the timecourse of either CAP 1 or 2 (a CAP, anti-CAP pair) was sufficient to eliminate inter-group differences in cortico-hippocampal rsfMRI connectivity observed with seed-based correlation analyses (p > 0.6, repeated measures two-way ANOVA, genotype-effect, Fig. 6E). Regression of all the other individual states, or the GS, did not substantially affect somatosensory-hippocampal rsfMRI coupling (p < 0.0005, all remaining CAPs and GS, repeated measures two-way ANOVA, genotype-effect, Fig. 6E). These results demonstrate that rsfMRI disconnectivity observed in *Chd8* mutants can be explained by the involvement of brain states characterized by aberrant co-activation topographies, opening the way to a interpretative reframing of rsfMRI disconnectivity in terms of non-canonical state engagement.

## Discussion

Here we describe time-varying patterns of spontaneous brain activity in terms of simple dynamical rules. We show that brain-wide patterns of fMRI activity can be classified into recurring states exhibiting coupled oscillatory activity. We further describe the temporal structure of these network transitions, and show that they occur at specific phases of global fMRI signal fluctuations, acting as coupled oscillators. We finally show that patterns of aberrant connectivity relevant for autism are associated with altered state topography and oscillatory dynamics.

Dynamic characterizations of rsfMRI signal via sliding window correlation have led to the description of spontaneous brain activity as a non-stationary phenomenon (Allen et al., 2012; Hutchison et al., 2013b; Grandjean et al., 2017). Our approach expands these investigations by revealing a set of fundamental principles guiding the spatiotemporal structure of resting state fMRI activity, implicating oscillatory transitions of recurring functional states as a fundamental level of organization of intrinsic brain activity. Importantly, the use of raw BOLD fMRI signal as the basis for the employed fMRI frame clustering provides an easily-interpretable and physiologically-relevant readout for describing time-varying patterns of spontaneous brain activity in non-correlative terms, and without computational constraints requiring the use of brain parcellations. Departing from other CAP-based approaches (Liu et al., 2013; Karahanoğlu and Van De Ville, 2015), we also introduce a series of empirical criteria to narrow down the selection of cluster numbers to increase the generalizability of our findings.

The identified brain states exhibit a composite spatial structure, providing novel insights into the macroscale functional organization of spontaneous network dynamics in the mouse brain. The presence of opposing DMN and LCN co-activation recapitulates a cardinal feature of human DMN organization (Fox et al., 2005), and supports the presence of a tight inverse coupling between these two neocortical systems (Popa et al., 2009; Gozzi and Schwarz, 2016). Interestingly, autoregressive pattern-finding algorithms have recently revealed a similarly competing relationship between DMN regions and constituents of the human task-positive network (TPN) (Yousefi et al., 2018), implicating the mouse LCN as a putative rodent homologue of the human TPN (Sforazzini et al., 2014). Importantly, the same states also provide a spatial delineation of the rodent DMN in non-correlative terms. The observed topography fully supports initial descriptions of this network to comprise in rodents thalamo-frontal as well as temporal associative and peri-hippocampal components (Sforazzini et al., 2014; Gozzi and Schwarz, 2016). The presence of opposing hippocampal and DMN activity in CAPs 5 and 6 is consistent with prior research implicating a functional interplay between these network systems in primates (Logothetis et al., 2012; Kaplan et al., 2016)

The observation of two global states characterized by widespread neocortical co-activation/de-activation provides key mechanistic clues as to the neural determinants of the oscillating states we described in this work. Optical imaging in awake and lightly anesthetized mice have revealed that spontaneous neural activity entails slow (< 0.1 HZ) neural waves spanning the entire cortex, as well as transient co-activations within neuro-anatomically constrained patterns of activity (Mohajerani et al., 2010; Matsui et al., 2016; Vanni et al., 2017). Concurrent hemodynamic and neural measurements have convincingly linked the two phenomena, demonstrating that transient co-activations embedded in global waves are converted into hemodynamic responses travel across homotopic dorso-cortical areas (Matsui et al., 2016; Schwalm et al., 2017). This relationship has been expanded to relate cortical co-activation patterns of calcium activity with spatially-structured hemodynamic fluctuations (Matsui et al., 2017), revealing recurring dorso-cortical CAPs that remarkably reproduce the contrasting involvement of DMN and LCN areas we described here (supplementary movies 1-2, 5-6). These spatial correspondences are consistent with a neural origin of the identified fMRI states, and support a view in which brain-wide fMRI fluctuations are inherently guided by intrinsic oscillatory cycling between slowly propagating neural activity. Non-conventional analyses of human rsfMRI network dynamics support this hypothesis. For example, five recurrent network configurations have been recently described using phase coherence as instantaneous connectivity metric using a regional parcellation of the human brain (Cabral et al., 2017). Similarly, data-driven approaches have uncovered putative temporal sequences of fMRI activity defined as lag threads of propagated fMRI signal in the human brain (Mitra et al., 2015) as well as two dominant, alternating quasi-periodic patterns involving DMN and TPN regions, reminiscent of our states 1 and 2 (Majeed et al., 2011; Belloy et al., 2018; Yousefi et al., 2018). Finally, whole-brain fMRI decomposition of human rsfMRI signal into CAPs produced a set of states that, although not recognized by the authors of the work as such, exhibit mirroring spatial configurations consistent with the oscillatory dynamics we describe here (see Figure S3 in Liu et al., 2013).

It should be emphasized that the light sedation regimen employed in our mouse rsfMRI datasets preserves cortical responsivity, without producing slow-waves or cortical hyper-synchronization (Orth et al., 2006; Gozzi et al., 2012). Moreover, this regimen is associated with high network specificity, and preserves thalamo-frontal fMRI connectivity (Sforazzini et al., 2014), a functional signature that is representative of conscious states (Liang et al., 2013). These lines of evidence suggest that the states we describe here and their oscillatory dynamics are representative of the repertoire of network configurations occurring during quiet wakefulness in the resting rodent brain. In keeping with this notion, *seed-based* CAP mapping of prefrontal and somatosensory regions in awake, restrained rats, revealed three non-redundant states, exhibiting remarkable spatial correspondence with our CAPs 2, 4, and 6, respectively (Liang et al., 2015).

A wave of recent investigations have linked fluctuations of the fMRI GS to vigilance (Wong et al., 2013), glucose metabolism (Thompson et al., 2016) and arousal mediated by ascending nuclei (Liu et al., 2018; Turchi et al., 2018). Our results are consistent with the hypothesis that the GS (when devoid of prominent artefactual contributions) encodes for key neuronal-relevant information, and support a view, in which each GS cycle is the sum of different, partially overlapping, network configurations. While the intrinsic drivers of these reconfigurations remain elusive, the observation of foci of co-deactivation in basal forebrain areas in CAP 3 (Fig. 3) recapitulates similarly contrasting patterns of global activity in cortical and basal forebrain regions observed in humans (Liu et al., 2018), and is consistent with a role of ascending modulatory activity in driving these oscillations (Turchi et al., 2018). The combined use of cell-type specific manipulations and rsfMRI (Giorgi et al., 2017) may permit to probe this mechanistic hypothesis, by enabling causal manipulations of ascending neurotransmitter systems.

Finally, the observation of altered state topography and oscillatory dynamics in a mouse line harboring an autism-associated mutation provides a novel interpretative framework for the description and interpretation of dysfunctional connectivity in brain disorders. Altered steady-state rsfMRI connectivity has been widely document in autism (Di Martino et al., 2013), and can be effectively recapitulated in mouse lines harboring mutations in ASD risk genes (Sforazzini et al., 2016; Liska et al., 2017; Michetti et al., 2017; Bertero et al., 2018). Our findings suggest that aberrant rsfMRI functional coupling reflect non-canonical patterns of regional co-activation and impaired state dynamics, thus providing a spatio-temporal description of brain dysfunction that can be used to relate basic neurophysiological mechanism (e.g. defective neural co-activation) to the deficient inter-regional communication observed in brain connectopathies. Notably, the statistical significance of the observed inter-strain differences appear to greatly exceed the sensitivity of conventional steady-state rsfMRI mapping. This finding opens the way to the use of multi-dimensional spatio-temporal mapping as a possible, clinically relevant criterion for patient stratification. Further investigation of the neural basis of canonical and altered state dynamics are warranted to pinpoint the exact neural drivers of these oscillating states and their aberrant topographies in models of human pathology.

In summary, our work documents a set of recurring oscillating brain states that govern the spatio-temporal organization of intrinsic brain function in the mammalian brain, allowing to describe time-varying patterns of spontaneous brain activity in terms of simple dynamical rules. These findings points at brain-wide oscillatory network activity as a fundamental level of organization of spontaneous brain activity, and add a novel interpretative dimension to the investigation of spontaneous brain activity, and its breakdown in brain disorders.

## Methods

### Ethical Statement

All in vivo experiments were conducted in accordance with the Italian law (DL 26/214, EU 63/2010, Ministero della Sanità, Roma) and the recommendations in the Guide for the Care and Use of Laboratory Animals of the National Institutes of Health. Animal research protocols were reviewed and consented by the animal care committee of the Istituto Italiano di Tecnologia. All surgical procedures were performed under anesthesia.

### rsfMRI data acquisition

Experiments were performed on a set of n = 40 adult (12-18 week-old) male C57Bl6/J mice. This is the main dataset that we have used throughout our article to describe rsfMRI dynamics, and all results presented below have been generated with this set of rsfMRI images, unless otherwise stated. The animal preparation protocol for experimental measurements has been described in great detail (Ferrari et al., 2012; Sforazzini et al., 2016). Briefly, mice were anaesthetized with isoflurane (5% induction), intubated and artificially ventilated (2%, surgery). The left femoral artery was cannulated for continuous blood pressure monitoring and terminal arterial blood sampling. At the end of surgery, isoflurane was discontinued and substituted with halothane (0.75%). Functional data acquisition commenced 45 min after isoflurane cessation. Mean arterial blood pressure was recorded throughout imaging sessions. Arterial blood gases (p_a_CO_2_ and p_a_O_2_) were measured at the end of the functional time series to exclude non-physiological conditions.

MRI data were acquired with a 7.0 Tesla MRI scanner (Bruker Biospin, Ettlingen) as previously described (Liska et al., 2015a), using a 72 mm birdcage transmit coil, and a four-channel solenoid coil for signal reception. For each session, high-resolution anatomical images were acquired with a fast spin echo sequence (repetition time (TR)/echo time (TE) 1200/15 ms, matrix 192 × 192, field of view 2 × 2 cm2, 18 coronal slices, slice thickness 0.60 mm). Co-centered single-shot blood-oxygen level dependent (BOLD) EPI time series were acquired using an echo planar imaging sequence with the following parameters: TR/TE 1200/15 ms, flip angle 30°, matrix 100 × 100, field of view 2 × 2 cm2, 18 coronal slices, slice thickness 0.50 mm, 500 (n= 21) or 1500 (n = 19) volumes and a total rsfMRI acquisition time of 10 or 30 minutes, respectively. All the group analyses were carried out on the first 500 timepoints (10 min). The single subject CAP analysis was limited to the n = 19 subjects in which we acquired 1500 timepoints.

To corroborate the reproducibility of our findings across independent datasets, we applied the whole analytical pipeline to an additional dataset composed of n = 41 male C57Bl6/J mice in which we acquired rsfMRI timeseries (n = 300, 6 min) using the same sedation protocol and image parameters employed in the present study. A characterization of functional network organization in this set of animals has been previously described (Liska et al., 2015). We refer to this animal cohort as to dataset 2. Finally, to assess the ability of our analytical pipeline to detect aberrant states in mouse models of brain pathology, we applied our analytical framework to a cohort of Chd8 haploinsufficient mice, a relevant subtype of autism spectrum disorder, which has been previously described to present aberrant rsfMRI network activity (Suetterlin et al., 2018). Briefly, rsfMRI imaging was performed on 15-18 week old mice (n=23 *Chd8*^+/+^; n=19 *Chd8*^+/−^, both lines have a C57Bl6/J background), each with 500 volumes (10 min), using the same animal preparation protocol and rsfMRI acquisition parameters of dataset 1. The control group in this third study was employed as a third independent dataset for the validation of our clustering procedure (dataset 3).

### Data pre-processing

Data preprocessing was carried out as recently described (Liska et al., 2017). Briefly, fMRI time series were despiked, motion corrected, and spatially normalized to an in-house mouse brain template (Sforazzini et al., 2014) yielding a final normalized spatial resolution of 0.1 × 0.1 × 0.5 mm^3^ (192 × 192 × 24 matrix). Head motion traces and the mean ventricular signal (average fMRI time series within a manually-drawn ventricle mask from the template) were regressed out. The resulting images were spatially smoothed using a Gaussian kernel of 0.5 mm FWHM, band-pass filtered using a 0.01 – 0.1 Hz band, and z-scored voxel-wise.

### Seed-based CAP analysis and spatial correspondence with rsfMRI correlation networks

To probe the relationship between networks inferred from conventional rsfMRI seed-based correlation analyses, and those described by high regional fMRI activity at only a few critical time points (Tagliazucchi et al., 2012; Liu and Duyn, 2013), we spatially averaged individual rsfMRI volumes (here referred to as “frames”) exhibiting peaks of regional activity in a set of *a priori* regions of interest. The employed averaging yields spatial maps of averaged spontaneous fMRI activity, which we termed seed-based mean CAPs. Seed location was chosen based on prior rsfMRI mapping in the mouse (Sforazzini et al., 2014; Liska et al., 2015). For each of the probed regions, we also computed a canonical group-level correlation map using the corresponding regional rsfMRI signal as seed. We next computed the spatial correlation between the seed-based CAPs and their corresponding correlation maps by retaining rsfMRI frames exceeding a predefined intensity threshold, covering the whole 0-99^th^ percentile range as previously described (Liu and Duyn, 2013). For illustrative purposes, we generated representative seed-based CAPs by averaging all the fMRI frames with the highest 15% BOLD signal intensity across all subjects (68 ± 5 out of 500 frames, mean ± SD, all subjects), and using a T threshold of 7, corresponding to p < 0.01, Bonferroni corrected (Liu and Duyn, 2013; Amico et al., 2014). For each region we also generated a group-level canonical seed-based correlation map, which we thresholded at T = 7, corresponding to p < 0.01, Bonferroni corrected).

### Whole-brain CAP analysis

We used spatial clustering of individual fMRI frames to identify whole-brain patterns of simultaneous co-activation of brain activity with voxel-resolution. Following prior studies (Liu et al., 2013; Liu and Duyn, 2013; Karahanoğlu and Van De Ville, 2015), we carried out frame-wise clustering of brain-wide mouse rsfMRI images by retaining the voxels that were in the top 10% or in the bottom 5% of all BOLD signal in a single time frame, a procedure employed to minimize spurious influence of random, non-physiological signal fluctuations in fMRI timeseries (Liu et al., 2013). The preprocessed fMRI frames were formatted into N-dimensional vectors (*t*_1_, *t*_2_, …, *t*_T_) with *T* being the number of frames, and *N* the amount of voxels. Such frames were clustered using the k-means++ algorithm (Arthur and Vassilvitskii, 2009), which partitions the vector-set into k clusters *C* = (*C*_1_, *C*_2_, …, *C*_k_) such that the sum of within-cluster distances D in the following equation is minimized:

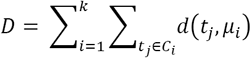

being μ_i_ the mean of the fMRI frames in each cluster C_i_, and d(t, μ_i_) the distance between the fMRI frame at time *i* and the cluster mean, measured as one minus the spatial Pearson’s correlation coefficient. The k-means++ algorithm provides an optimized initialization of each centroid (seeding), such that distant centroids have a higher chance of being chosen as starting seeds for the clustering procedure. The addition of this step has been shown to outperform conventional k-means with random seeding both in terms of clustering performance and speed (Arthur and Vassilvitskii, 2007).

We note here that the above described intensity masking does not affect the final results, and operating the clustering, as well as all the successive phase analyses, directly on unmasked fMRI data did not change the results qualitatively, either in terms of CAP spatial shape or in terms of phase results. Indeed, clustering the unmasked data led only to a marginal change (fraction 1.64%) in the time frames that were assigned to different clusters by either the masked or the unmasked procedure, and the spatial correlation between the CAPs obtained with or without masking was higher than > 0.98 for all the states.

k-means clustering was carried out on the whole rsfMRI dataset 1 (40 × 500 frames), with k ranging from k = 2 up to 20, using Pearson correlation between fMRI data in different time frames as clustering distance measure, and using 15 replicates with 500 iterations each. Departing from previous investigations in which the number of cluster was selected ad hoc (Liu et al., 2013; Liu and Duyn, 2013; Karahanoğlu and Van De Ville, 2015), we carried out k number selection generalizing a more quantitative procedure, aimed at identifying the most robustly occurring co-activation patterns, which was previously employed to map robust, recurrent states in electrophysiological recordings (Logothetis et al., 2012). Briefly, we first computed, for increasing k, how much variance is explained by the clustering algorithm (Fig. S1B), defined as the ratio between the between-cluster variance and the total variance (within-cluster + between-cluster variance). Within-cluster variance was computed as the averaged (over clusters) sum of square distances between elements in a cluster and its centroid. Between-cluster variance was computed as the averaged square distance between a cluster centroid and the centroid of all clusters or centroid of all data (Goutte et al., 1999). Higher values of explained variance correspond to a better description of the dataset, and that optimal k values can be identified in the “elbow” region of the explained variance plot, after which only marginal variance is gained by further refining the partition. We thus selected the value k as the largest value within the elbow region that insured full reproducibility of the corresponding CAPs across the three independently collected datasets used in this study. To this purpose, we progressively increased k until *(a)* the dataset variance explained by k clusters was larger than the variance explained by k-1 clusters and *(b)* a replication of the clustering procedure on two additional independent rsfMRI datasets (n = 41 and n = 23, respectively), would result in anatomically equivalent co-activation patterns (defined as between dataset CAP correlation > 0.45, which corresponds to a highly significant correlation, p <10^−5^, permutation test, Fig. S1). This procedure identified k = 6 CAPs conserved across all the three rsfMRI datasets (Fig. 3 and Fig. S1). To compute the centroid of each cluster (which we took as a CAP), the fMRI frames assigned to each cluster were averaged voxel-wise, and normalized to T-scores (p < 0.01, Bonferroni corrected), permitting to visualize the mean voxel-wise distribution of fMRI BOLD signal for each of the identified CAP. For visualization purposes, the obtained maps were thresholded to T-scores > 7, corresponding to a p<0.01, Bonferroni corrected. For each CAP we next computed its occurrence rate (i.e. the proportion of frames assigned to each CAP) and duration (i.e. the average number of consecutive frames belonging to the same CAP at each occurrence) for each of the n = 40 subjects. We also computed between-CAP spatial similarity, defined as pair-wise spatial correlation between all the identified CAPs.

To assess the ability of our approach to detect CAPs also at the subject level, we repeated the clustering analysis in a subset of 19 subjects of the main dataset for which we acquired extended (30 min, 1500 frames) time series, using the group-level CAPs as initial centroids. The CAPs found in each subject were next matched to the group-level templates using the Hungarian Algorithm (Kuhn et al., 1955), and their spatial correlation was computed. To illustrate CAP incidence across subjects, we mapped, for each voxel in a CAP, the proportion of subjects which had a significant co-activation (T-test, p < 0.05, FDR corrected) and equal co-activation sign (i.e. positive or negative BOLD signal) of the corresponding group-level CAP template.

### CAP dynamics

To investigate the dynamics of CAP evolution, we generated a CAP-to-frame correlation time course (which we term “CAP time course”). We computed these spatial correlations in each time frame as the Pearson correlation between the centroid of the considered cluster and the masked fMRI activity in the considered time frame of a given subject as previously described (Liang et al., 2015). For plotting purposes, we normalized the CAP time courses into SD units. This normalization does not affect in any way either the power spectra profile or the distribution of the phase values. To describe the assembly and disassembly dynamics of CAPs, we used the method devised by Liang et al. (2015). Briefly, for each CAP we selected the fMRI frames corresponding to the local maxima within the CAP’s time course, limiting the selection of peak events to 2% of all the concatenated frames, and adjusting this number for each CAP’s occurrence rate. For example, for CAP 1 (occurrence rate = 0.18) we sampled 0.18 × 400 frames corresponding to the CAP’s time course local maxima. Each selected frame was set as a time *t* = 0 reference event, and we next sampled frames within a −30 ≤ *t* ≤ 30 repetition interval. All events were time-lock averaged, leading to a dynamic portrayal of mean temporal evolution of CAP assembly and disassembly in the form of a concatenated frames (supplementary movies S1-6).

The temporal structure of the CAP time courses for each subject was also assessed by computing its power spectrum. To compute the instantaneous phase of the GS in the infra-slow oscillation range, we first band-pass filtered the GS time-courses between 0.01-0.03 Hz. We then used the Hilbert Transform (Montemurro et al., 2008) to decompose the GS signal into an analytical signal with a characteristic instantaneous phase and amplitude. We divided each subject’s instantaneous GS phase signal into cycles in the range [0, 2π], and within each cycle, collected the GS phase values at each CAP’s occurrence. Using the same filter design described above, we filtered and again normalized the CAP time courses, and sampled the GS phase at each CAP occurrence only when the CAP time course at that instant was above 1 SD, in order to ensure that a specific frame pertained to a specific CAP. Using the Matlab CircStats toolbox (Berens et al., 2009), we computed circular statistics of the obtained distribution of GS phases at each CAP, and represented their dispersion in a cosine cycle representing a GS oscillation, using the circular variance to quantify dispersion around the circular mean. To probe the presence of phase-coupling between CAP occurrence and GS oscillatory dynamics, we computed the angular differences (phase differences) of the GS between occurrences of a given CAP within a GS cycle, and occurrences of another CAP within the previous, current, and subsequent cycle. Again, GS phase samples at each CAP were only considered if their filtered values at that instant were above 1 SD at the corresponding instances.

### CAP dynamics in a genetic model of autism

The same analytical pipeline described above was applied to rsfMRI timeseries recorded in n = 23 *Chd8*^+/+^ and n=19 *Chd8*^+/−^ littermates (Suetterlin et al., 2018) after standard preprocessing step. CAPs were computed independently in each group. We used k = 6, as this value ensured highest cross-dataset reproducibility of the identified clusters as described in the Results section (Fig. S1). Inter-group CAP maps were matched using the Hungarian algorithm according to their spatial similarity. Inter-group differences in CAP anatomy were mapped using a two-sample t-test and family-wise error (FWE) cluster correction (p < 0.05, cluster-defining threshold T(40) = 2.8). We then computed CAP features and CAP time-courses for all subjects in each strain, and sampled the filtered GS-phase distributions at each CAP occurrence for each group of mice independently. Differences between preferred GS-phase at CAP occurrences were computed using a William-Watson test for circular mean homogeneity, Bonferroni corrected for six comparisons. We also sampled the angular differences (phase differences) of the GS between occurrences of a given CAP within a GS cycle, and occurrences of another CAP within the previous, current, and subsequent cycle. Again, GS phase samples at each CAP were only considered if their filtered values at that instant were above 1 SD at the corresponding instances.

Finally, to assess the involvement of individual CAPs in the seed-based rsfMRI correlation differences previously described between hippocampal and motor-sensory areas (Suetterlin et al., 2018), we computed differences in whole-brain correlation maps between a 3 × 3 × 1 voxel seed placed bilaterally in the somato-sensory cortex. Inter-group differences were assessed using a two-sample t-test and family-wise error (FWE) cluster correction (p < 0.05, cluster-defining threshold T(40) = 2.8). We then recomputed Pearson correlation between this seed and selected foci of over-connectivity in the hippocampus. The rsfMRI correlation profiles were computed before and after regressing each CAP’s time-course independently in each subject. Differences in rsfMRI correlations were assessed by means of a repeated measures two-way ANOVA.

## Acknowledgments

A.G. acknowledges funding by the Simons Foundation (SFARI 400101, A. Gozzi) and the Brain and Behavior Foundation (2017 NARSAD, Independent Investigator Grant 25861). M.A.B was supported by grants from the Simons Foundation (SFARI 344763) and Medical Research Council (MR/K022377/1).

## Supplementary Figures

**Figure S1.**
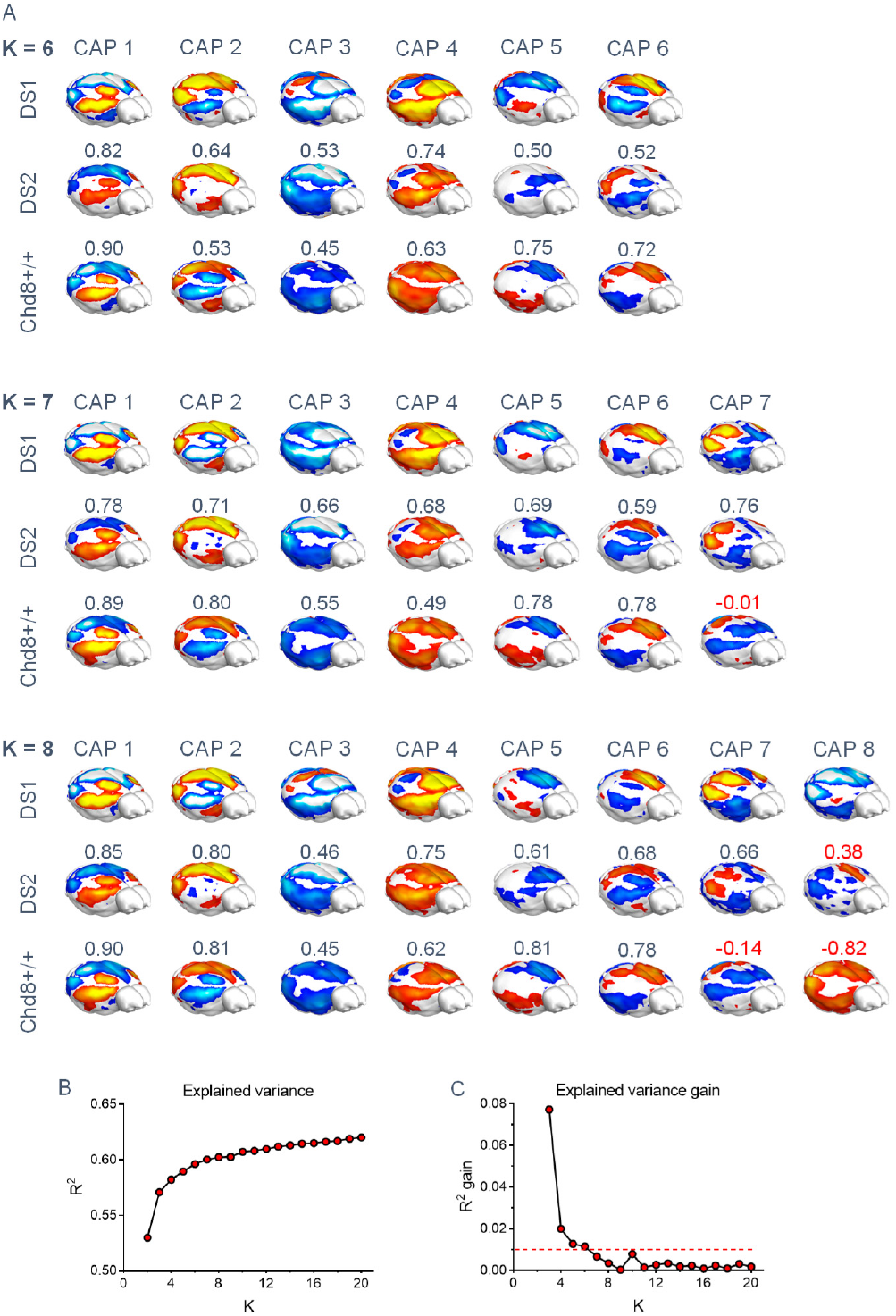
Selection of optimal number of clusters. (A) Whole-brain representations of CAPs found with k = 6, 7, and 8 in dataset 1 (n = 40, 500 fMRI frames per subject), and their matched CAPs found in dataset 2 (n = 41, 300 frames per subject) or dataset 3 (CHD8^+/+^ wildtype mice, n = 23, 450 frames per subject). Note that CAPs 1-6 are recurrently found in all datasets with k = 7 and 8, while additional CAPs are less reproducible across datasets. (B) Variance explained by clustering dataset 1 with k = 2 – 20. (C) Percentage gain in variance explained when advancing from k-1 to k.

**Figure S2.**
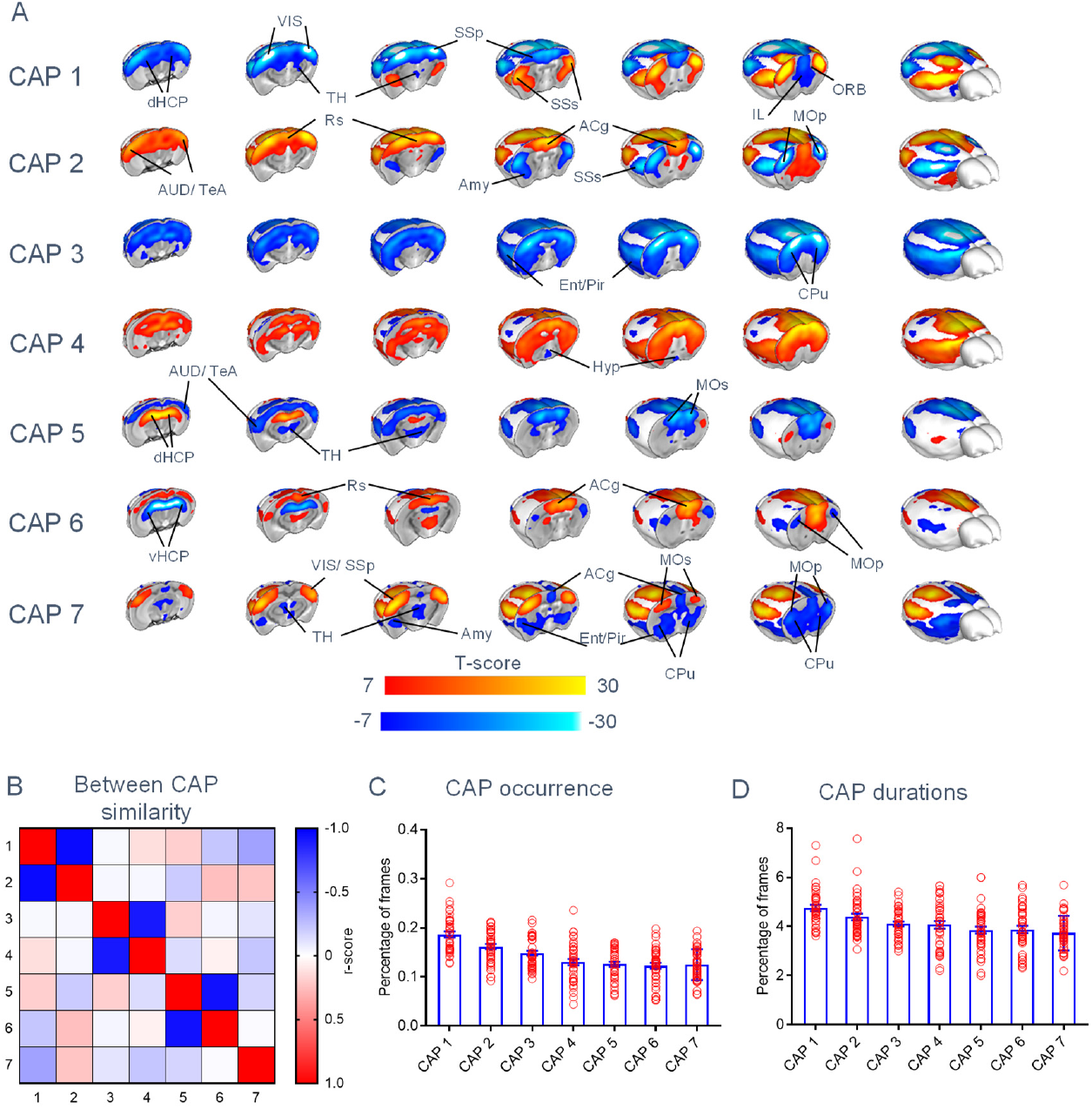
Brain states obtained with k = 7. (A) Whole brain representation of the 7 brain states (CAPs) we identified at the group level in dataset 1, using k = 7. Red/yellow indicates co-activation (i.e. high fMRI BOLD signal) while blue indicates co-deactivation (i.e. low fMRI BOLD signal) (p < 0.01, Bonferroni corrected). CAPs are organized according to their occurrence rates. Panels (C) and (D) illustrate CAP occurrence rate and mean duration, respectively (mean +/− SEM). Abbreviations: ACg – Anterior Cingulate cortex; AUD – Auditory cortex; dHC – dorsal Hippocampus; vHC – ventral Hippocampus; HT – Hypothalamus; ILA – Infralimbic Area; LAN – Lateral Amygdalar Nucleus; MOp – primary Motor cortex; Mos – secondary Motor cortex; ORB – Orbitofrontal cortex; PIR – Piriform Area; PL – Pallidum; Rs – Retrosplenial cortex; SSp – primary Somatosensory cortex; SSs – secondary Somatosensory cortex; ST – Striatum; TeA – Temporal Association cortex; TH – Thalamus; VIS – Visual cortex.

**Figure S3.**
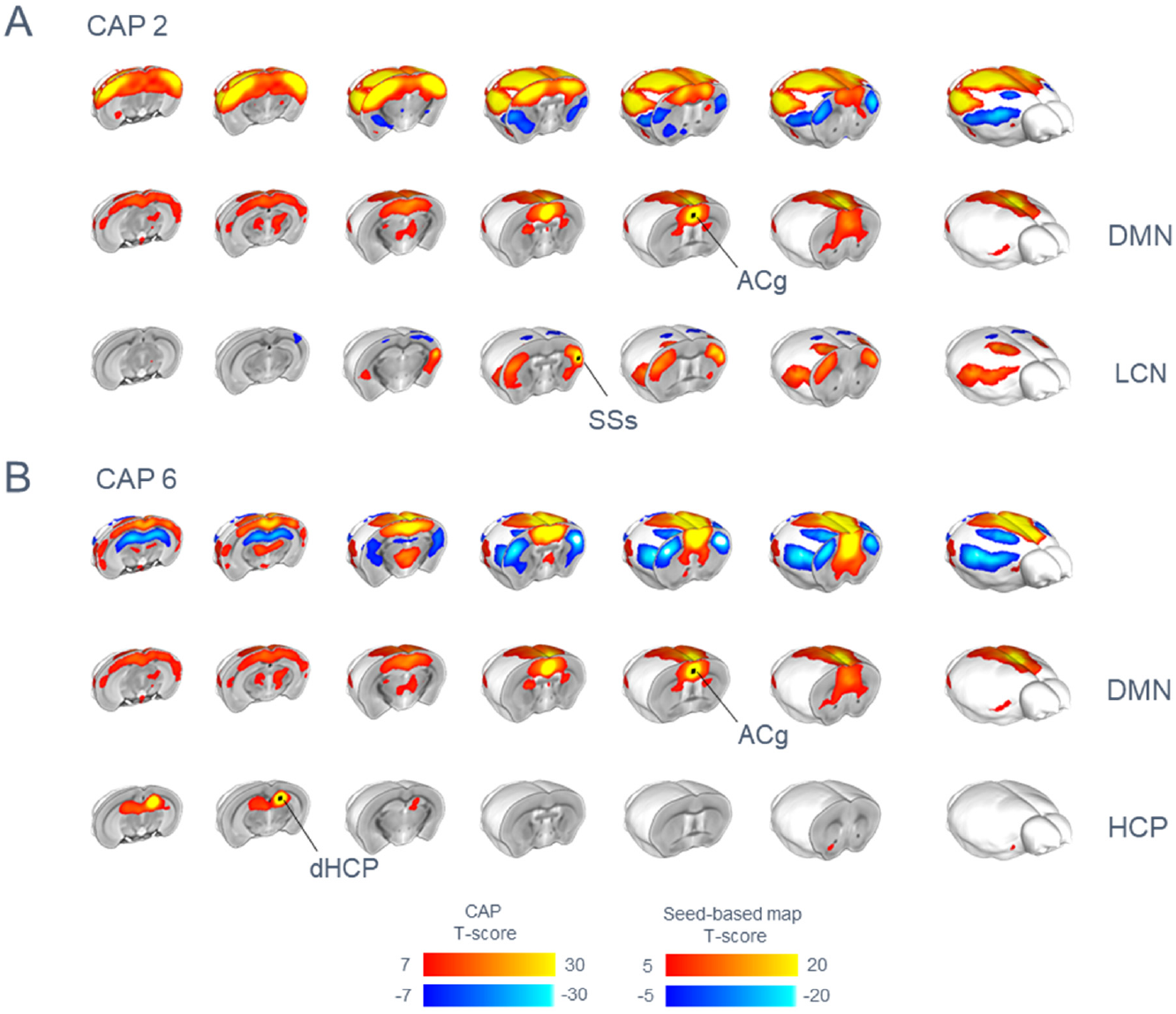
Brain functional states encompass known rsfMRI networks of the mouse brain. CAP spatial patterns (top row) can be decomposed into conventional rsfMRI network constituents generated with a seed based analysis (seed location indicated by black dots). (A) For example, CAP2 (top row) shows co-activation of regions of the mouse DMN and co-deactivation in the mouse latero-cortical networks (LCN). Similarly, CAP6 (B) exhibits contrasting involvement of regions of the DMN, and dorsal hippocampal network (HCP).

**Figure S4.**
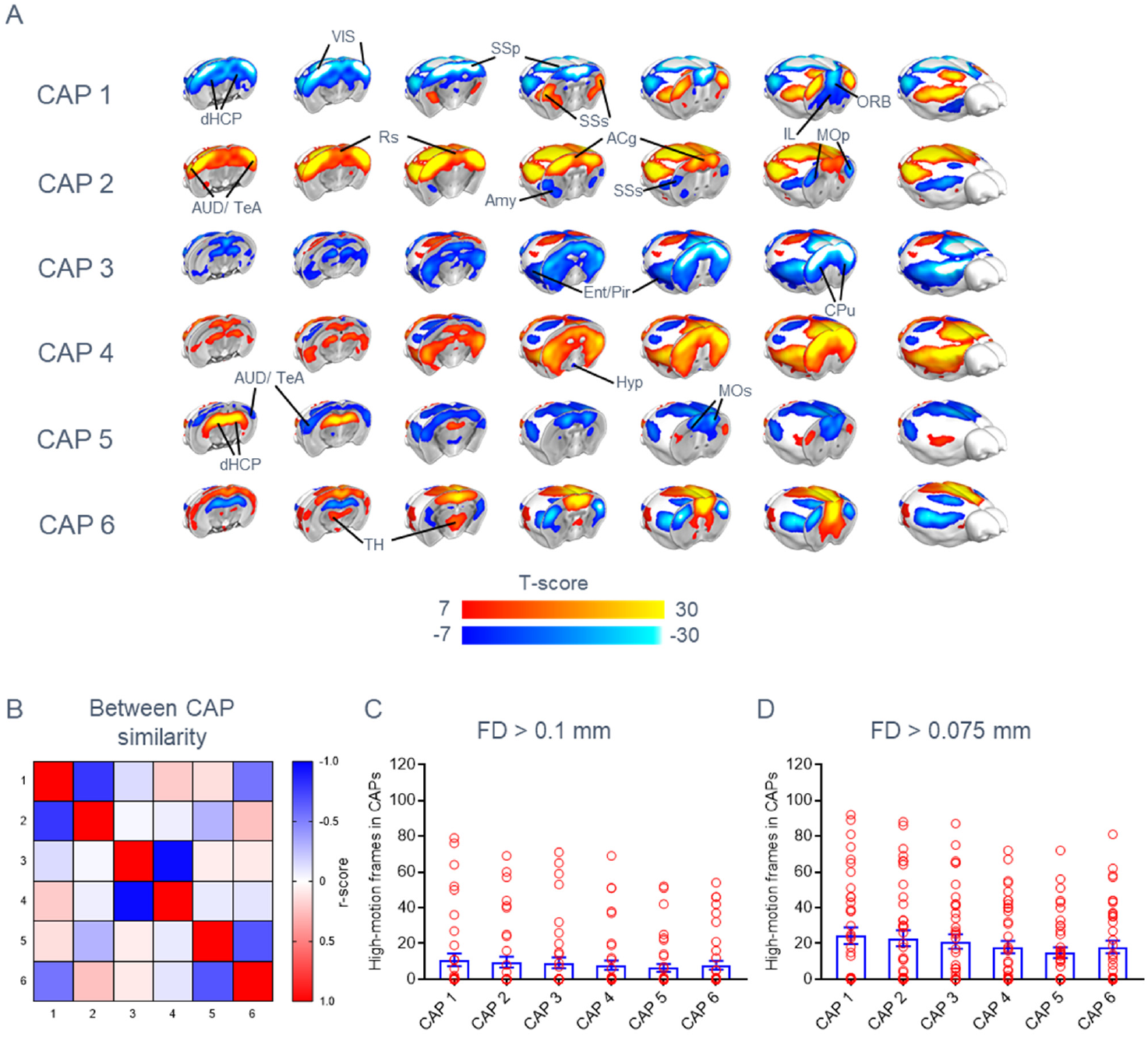
CAPs are not contaminated by motion. (A) CAPs obtained by clustering rsfMRI frames upon censoring of putative motion-contaminated frames using a frame-wise displacement (FD) threshold of 75 µm. (B) Between-CAP spatial similarly (correlation coefficient). (C-D). Distribution of putative motion-contaminated frames across CAPs at two strict FD thresholds (means +/− SEM).

**Figure S5.**
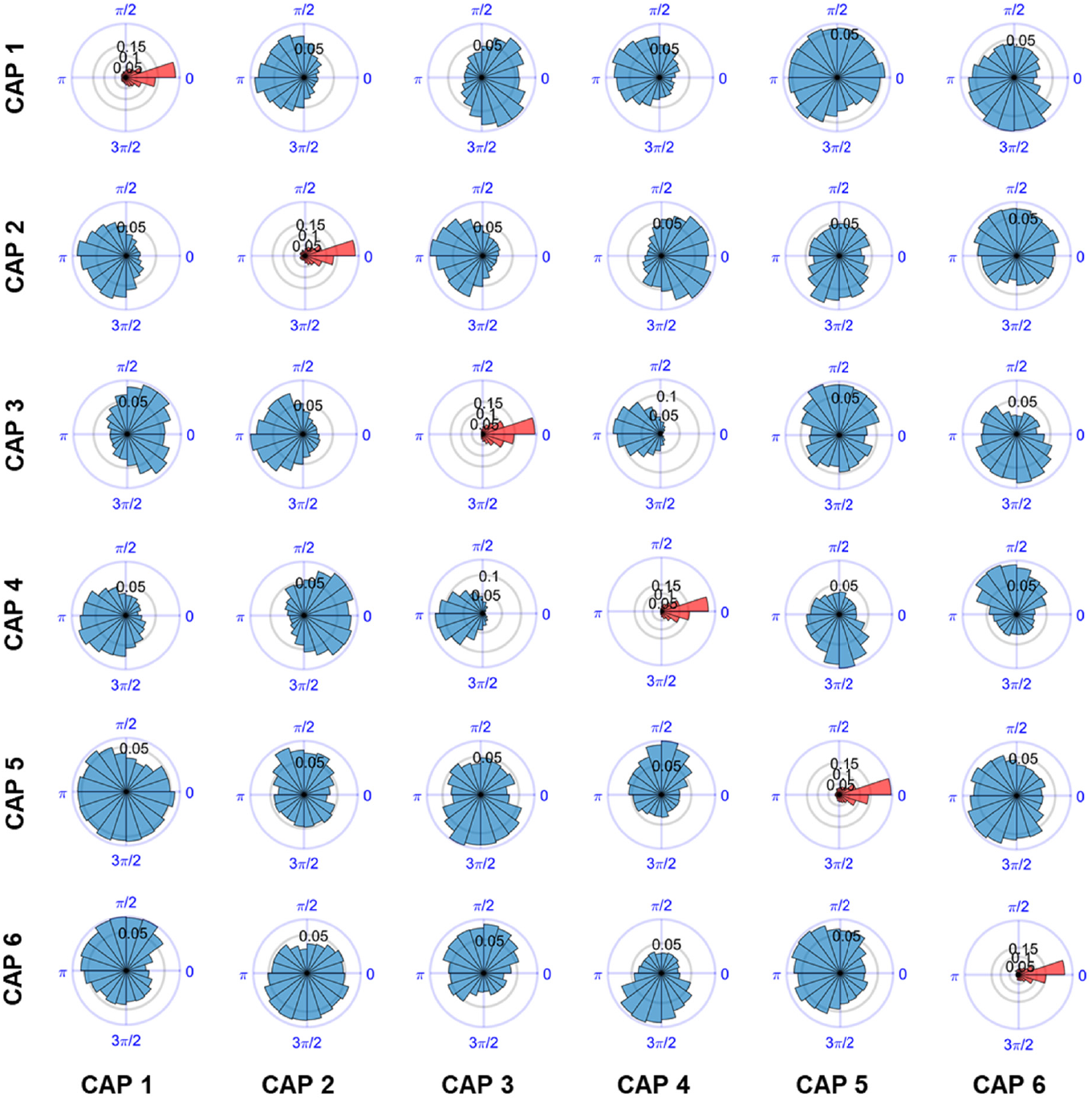
Functional states act as coupled oscillators. Global signal phase differences between CAP occurrences. Each panel corresponds to the circular distribution of GS phase differences between occurrences of a CAP inside a GS-cycle (rows), and the occurrences of another CAP within the same cycle and across GS cycles that were immediately adjacent in time (columns). All distributions significantly deviate from circular uniformity (Raleigh test, p < 0.05).

**Figure S6.**
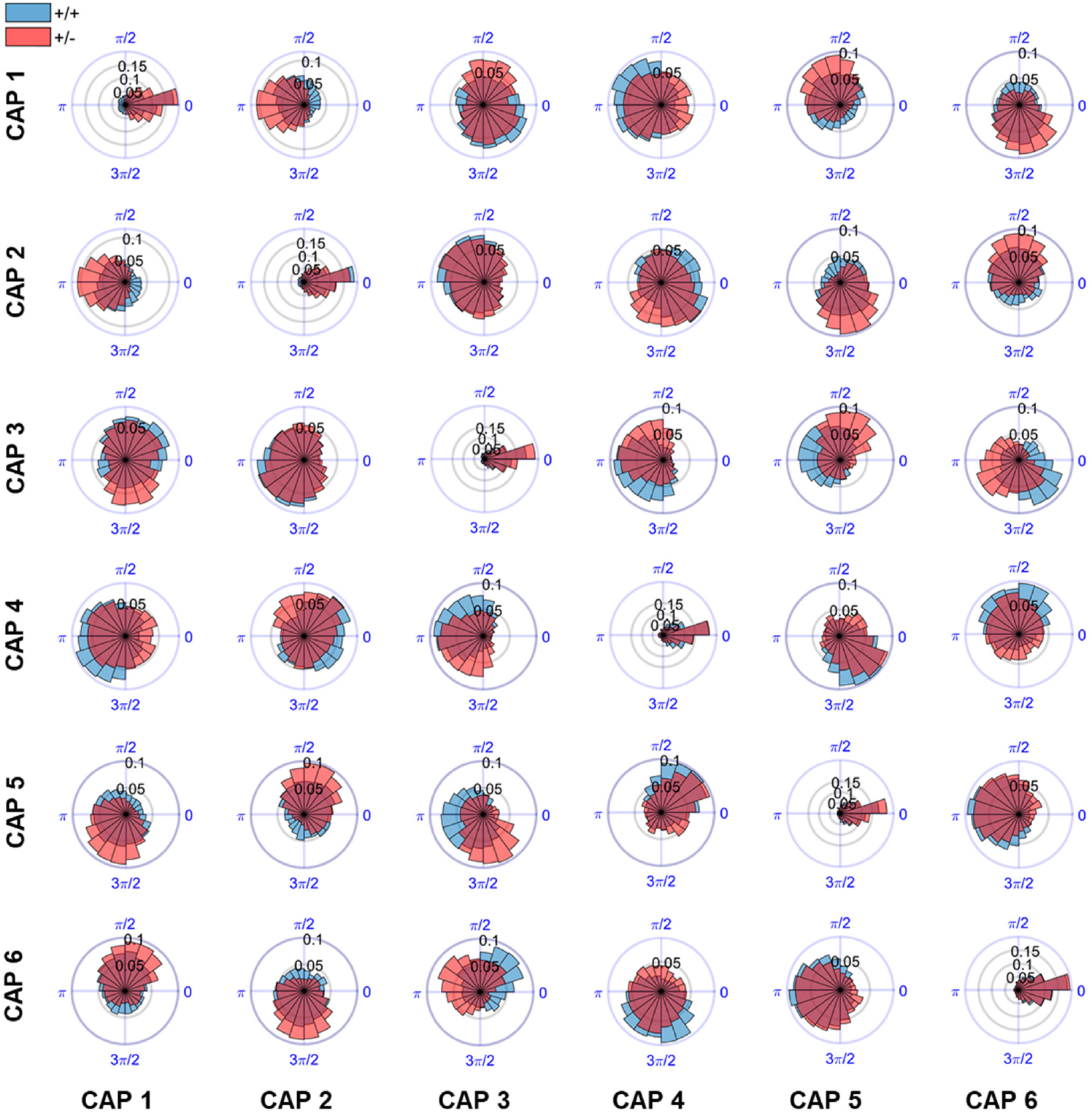
Altered GS phase difference between CAPs in a genetic model of autism. Each panel corresponds to the circular distribution of GS phase differences between occurrences of a CAP inside a GS-cycle (rows), and the occurrences of another CAP within the same cycle, and across GS cycles that were immediately adjacent in time (columns). All distributions significantly deviate from circular uniformity (Raleigh test, p < 0.05). Blue distributions correspond to CAPs from the control group (Chd8^+/+^), and red overlaid distributions correspond to CAPs from the mutant group (Chd8^+/−^). Black asterisks denote significant differences between circular means in each panel (William Watson test, p < 0.05).

